# Evaluating the risk of SARS-CoV-2 transmission to bats using a decision analytical framework

**DOI:** 10.1101/2021.05.28.446020

**Authors:** Jonathan D. Cook, Evan H. Campbell Grant, Jeremy T. H. Coleman, Jonathan M. Sleeman, Michael C. Runge

## Abstract

Preventing wildlife disease outbreaks is a priority issue for natural resource agencies, and management decisions can be urgent, especially in epidemic circumstances. With the emergence of SARS-CoV-2, wildlife agencies were concerned whether the activities they authorize might increase the risk of viral transmission from humans to North American bats but had a limited amount of time in which to make decisions. We provide a description of how decision analysis provides a powerful framework to analyze and re-analyze complex natural resource management problems as knowledge evolves. Coupled with expert judgment and avenues for the rapid release of information, risk assessment can provide timely scientific information for evolving decisions. In April 2020, the first rapid risk assessment was conducted to evaluate the risk of transmission of SARS-CoV-2 from humans to North American bats.

Based on the best available information, and relying heavily on formal expert judgment, the risk assessment found a small possibility of transmission during summer work activities. Following that assessment, additional knowledge and data emerged, such as bat viral challenge studies, that further elucidated the risks of human-to-bat transmission and culminated in a second risk assessment in the fall of 2020. We update the first SARS-CoV-2 risk assessment with new estimates of little brown bat (*Myotis lucifugus*) susceptibility and new management alternatives, using findings from the prior two risk assessments and other empirical studies. We highlight the strengths of decision analysis and expert judgment not only to frame decisions and produce useful science in a timely manner, but also to serve as a framework to reassess risk as understanding improves. For SARS-CoV-2 risk, new knowledge led to an 88% decrease in the median number of bats estimated to be infected per 1000 encountered when compared to earlier results. The use of facemasks during, or a negative COVID-19 test prior to, bat encounters further reduced those risks. Using a combination of decision analysis, expert judgment, rapid risk assessment, and efficient modes of information distribution, we provide timely science support to decision makers for summer bat work in North America.

---

The emergence of severe acute syndrome coronavirus 2 (“SARS-CoV-2”) occurred in the fall of 2019 and quickly presented immediate and apparent health risks to humans worldwide. By March 2021, the novel pathogen and associated coronavirus disease (“COVID-19”) had resulted in over 34 million documented human disease cases and over 800,000 deaths globally (Dong and Du 2020; Johns Hopkins COVID-19 Dashboard). While the human health risks of COVID-19 are clear, empirical information on the risk to wildlife is less available, and thus there remains concern among North American natural resource managers for the potential that SARS-CoV-2 could be transmitted to wildlife from infected humans. Bats are a group of primary focus following the detection of a closely related betacoronavirus in a horseshoe bat (*Rhinolophus affinis*) in eastern Asia (Olival et al. 2020); however, empirical study to directly assess the threat that SARS-CoV-2 presents to bats remains limited. Thus, there is an ongoing need for formal risk assessments that can best integrate existing knowledge and guide pressing management decisions regarding activities that require human-bat interaction.

When first confronted with the potential for SARS-CoV-2 exposure and infection in North American bats, natural resource managers had a limited suite of options to reduce the associated risk, including: proceeding as usual with minimal restrictions; placing a moratorium on all work under their authority that may elevate risk; or adopting mitigation actions thought to reduce risks. However, justification for selecting any one of these actions was a challenge because of the many uncertainties that obscured identification of an optimal approach. A few of the most pressing uncertainties surrounded bat species susceptibility, dominant transmission pathways, and estimation of the relative exposure and transmission risk that different human-bat interactions presented. Decisions had to be made without waiting for research that could reduce these uncertainties. To help managers make urgent decisions with limited information and under uncertainty, a series of rapid risk assessments were performed in 2020 using a decision-making approach that helped to: (1) identify agency objectives; (2) guide the development of quantitative models that were explicitly linked to agency objectives; (3) maximize the utility of available data and knowledge; and, (4) assess management alternatives under dynamic and frequently changing conditions.

In April 2020, a first assessment was completed that estimated the risk of SARS-CoV-2 transmission to North American bats during summer activities. The assessment was guided by a structured decision-making approach using an interagency team to evaluate the best course of action for management agencies based on the identified objectives, uncertainties, and a risk model that explicitly linked objectives and possible mitigation actions (Runge et al. 2020). The focal species for that assessment was the little brown bat (*Myotis lucifugus*) and risks associated with conducting research, survey, monitoring, and management (RSM), wildlife rehabilitation (WR), and wildlife control (WC) activities during the North American spring and summer seasons were assessed. The primary RSM activities of concern were those that put scientists in close proximity to bats as part of efforts to study and mitigate the effects of white-nose syndrome (“WNS”; e.g., Hoyt et al. 2019), a fungal disease that has caused declines of over 90% in affected little brown bat populations (Cheng et al. 2021); activities from other workers included removal and exclusion of bats from human dwellings (WC) and the care of injured bats (WR). Much of the information used for the assessment was derived from a formal process of expert judgment, as empirical data about the human and wildlife potential of SARS-CoV-2 were largely unknown at the time.

The first assessment by Runge et al. (2020) estimated that the risk of SARS-CoV-2 transmission from humans to bats was non-negligible and that the risk could be reduced if well-fitted N95 respirators (a type of mechanical filter capable of removing viral particles from exhaled breath of infectious individuals) and other protective clothing were used during all human-bat interactions. In the analysis, critical uncertainties remained–most notably a high level of uncertainty about the probability of bat susceptibility. The authors noted that the existence of a decision framework complete with objectives, a quantitative risk model, and management alternatives (e.g., mandating use of N95 respirators, prohibiting certain permitted activities) could be used to rapidly update decisions as more empirical information was gained.

In the fall of 2020, another assessment was conducted that evaluated the risk of human-to-bat transmission of SARS-CoV-2 during winter research activities. Winter activities primarily occur in enclosed spaces, such as hibernacula and winter roosts, that were thought to increase the potential risk of exposure of the bats to aerosolized virus emitted by scientists (Cook et al. 2021). Thus, included in that assessment were new data on the effectiveness of facemasks to reduce viral emission of infectious humans, and new knowledge regarding the susceptibility of bat species to SARS-CoV-2. Importantly, by the time of the second assessment, two bat challenge studies had been conducted in laboratory settings; one found no viable infection in big brown bats (*Epstesicus fuscus*), and another found that fruit bats (*Rousettus aegyptiacus*) were somewhat susceptible (Hall et al. 2020, Schlottau et al. 2020). Additionally, studies on species-specific angiotensin-converting enzyme 2(“ACE2”) sequences, an indicator of viral binding potential, shed further light on potential bat susceptibility (e.g., Damas et al. 2020). In aggregate, experts estimated a much lower, and less uncertain, probability of infection for several bat species, including the little brown bat (Cook et al. 2021). New information about the importance of aerosols for human disease transmission and the effectiveness of other personal protective equipment (PPE) and COVID-19 testing for preventing exposure was incorporated into a risk model for winter bat work.

As we transition into another summer season (2021), the empirical data on bat susceptibility, human viral shedding dynamics, and the potential effectiveness of facemasks and COVID-19 testing to reduce bat exposure have provided sufficient justification to revisit the initial decision analysis and summer risk assessment. In this paper, our objective was to provide updated risk estimates for summertime RSM, WC, and WR activities. We first confirm that the structural elements of the decision framing from Runge et al. (2020) remain relevant to agencies considering summer bat work. We then update probability of susceptibility estimates for the little brown bat based on Cook et al. (2021) and re-evaluate the risk of SARS-CoV-2 human-to-bat transmission during summertime RSM, WC, and WR activities. We evaluate the effectiveness of COVID-19 testing, in addition to several new types of enhanced PPE, for their ability to prevent exposure and mitigate risk. We highlight the strengths of decision analysis to organize, evaluate, and improve time-sensitive decisions using expert knowledge, strategic decision framing, and frequent updating. We also provide a brief synopsis of a few institutional hurdles that challenged the delivery of our risk assessments to decision-makers and provide some potential options to improve the speed of decision-relevant science in future studies.

## METHODS

### Decision Framing and General Approach

The initial decision framing for SARS-CoV-2 transmission risk from humans to bats formed the basis for the results presented in Runge et al. (2020) and Cook et al. (2021). A diverse group of state and federal decision makers were involved in the framing, and as a result, it captured many of the objectives and management alternatives that agency decision makers were considering at the time. The framing of a decision may change over time and can lead to different structuring of the problem and resulting models. Therefore, to update the summer risk assessment we first revisited the original decision framing for spring and summer work with the original guidance committee from Runge et al. (2020). During our meeting, agency participants indicated that the decision context and all objectives remained the same. Of particular relevance to this assessment were objectives related to:

1. minimizing the morbidity and mortality of wild North American bats resulting from infection with SARS-CoV-2 or from management actions meant to mitigate transmission,
2. minimizing the risk of SARS-CoV-2 becoming endemic in any North American bat population through sustained bat-to-bat transmission,
3. maintaining or maximizing the ability of WC and WR to carry out their functions for the benefit of humans and wildlife, and
4. maximizing the opportunities for scientific research on bats and within bat habitats.

For a complete summary of objectives, including the full text of the select objectives presented here, see Runge et al. (2020).

Based on the agreement in objectives between the first summer assessment and this study, the existing infection risk models developed by Runge et al. (2020) remained useful for estimating risk, but needed to be updated to include new and relevant information. Conceptually, new information from empirical study and experts can reduce critical uncertainties and can be incorporated into the existing Runge et al. (2020) framework as indicated in Figure 1 by the gray, dashed arrows. In the following sections, we describe the three infection risk models for RSM, WR, and WC activity types from the previous assessment and then revise them to include new information (blue highlighted steps of Figure 1: “New Information (monitoring, research)” and “Update Predictive Models (Learn)”). We then provide updated estimates on bat risk and mitigation in the results that can help to evaluate the consequences of SARS-CoV-2 risk management strategies (Figure 1: “Evaluate Consequences”).

**Figure 1.**
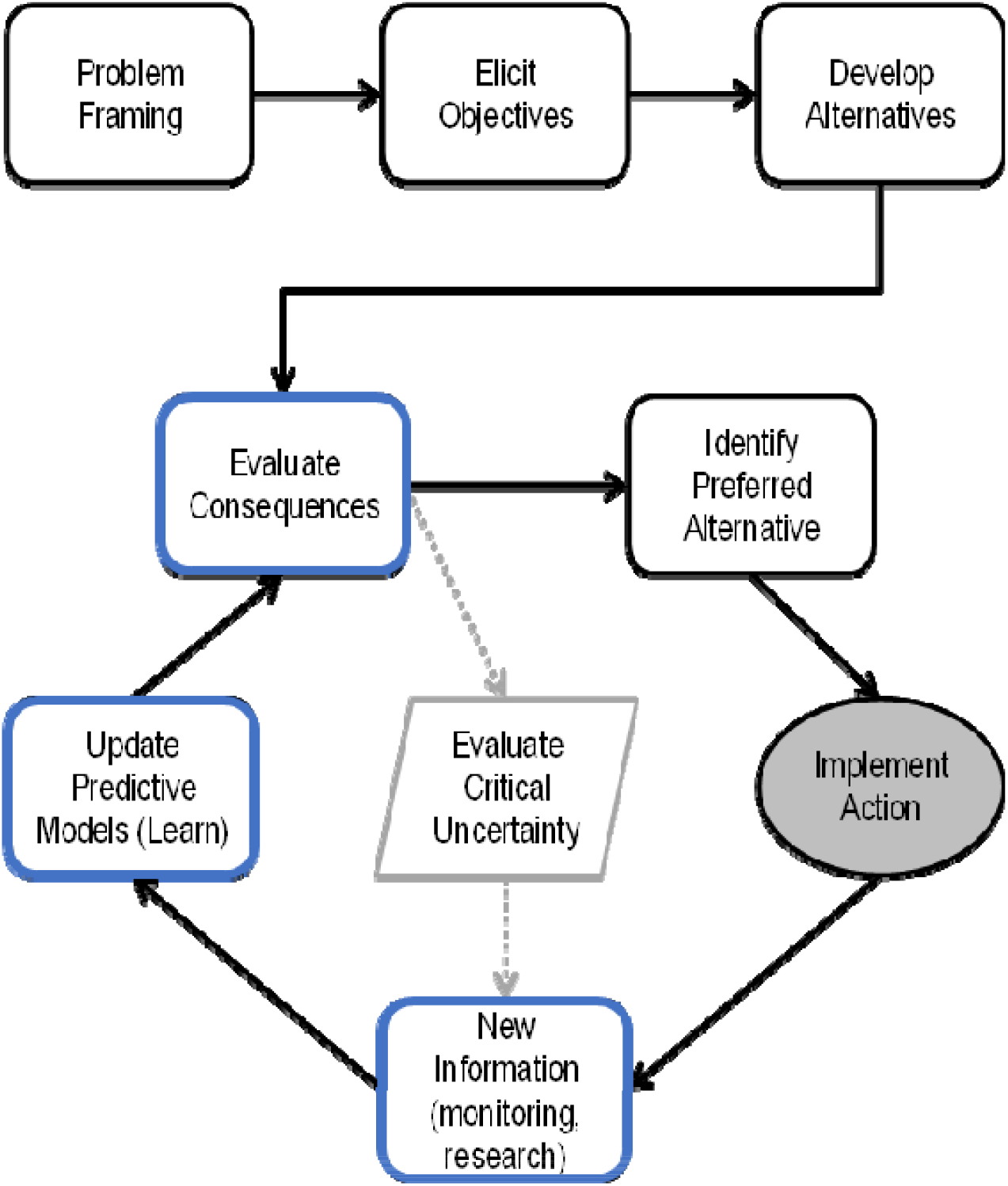
Steps of decision analysis, including the option to revisit consequences based on newly generated knowledge or data. In April 2020, Runge et al. (2020) worked with state and federal decision makers to frame the decision and produce risk estimates that were useful to guide management actions (Runge et al. 2020). Based on new knowledge and data that evaluated critical uncertainties from Runge et al. (2020) (gray dashed arrows and central gray outlined polygon), we revisited several steps (boxes outlined in blue) to rapidly re-evaluate the risk of SARS-CoV-2 transmission during summer RSM, WC, and WR activities. Frequently updating risk assessments using the best available science may help decision makers implement actions that best achieve management objectives (gray filled oval). Adapted from Runge (2011).

### RSM Infection Risk Model

The RSM infection risk model was calculated from three encounter types: workers handling bats, workers in proximity to bats in a shared enclosed space, and workers in close proximity to bats but not in a shared enclosed space. The expected number of infected bats resulting from research, survey, or monitoring activities is the sum of the expected number of bats infected through each of the three encounter types:

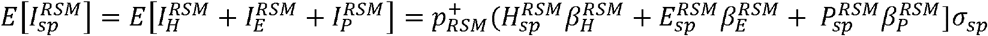

where

*I^RSM^* is the number of infected bats through each of three encounter pathways (*H* = handling of bats; *E* = exposed in a shared enclosed space; *P* = encountered not in an enclosed space)
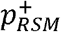 is the probability that someone conducting RSM work is actively shedding SARS-CoV-2 virus on any given day of the 2021 active season;
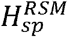 is the total number of bats handled during the 2021 active season;
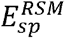 is the total number of bats exposed in a shared enclosed space, but not handled, during the 2021 active season;
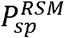 is the total number of bats encountered, but not in an enclosed space or handled, over the course of the 2021 active season;
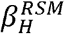 is the probability that a bat handled by a RSM scientist who was actively shedding virus would be exposed to the virus (an “exposure probability”) in the absence of any new restrictions, regulations, or protocols, taking into account the handling time typical of RSM activities;
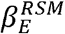 is the probability that a bat in an enclosed space within a 6-foot proximity of (but not handled by) a RSM scientist who was actively shedding virus would be exposed to the virus (an “exposure probability”) in the absence of any new restrictions, regulations, or protocols;
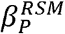 is the probability that a bat **not** in an enclosed space within a 6-foot proximity of (and not handled by) a RSM scientist who was actively shedding virus would be exposed to the virus (an “exposure probability”) in the absence of any new restrictions, regulations, or protocols; and
σ_*sp*_ is the species-specific probability that a bat exposed to a sufficient viral dose of SARS-CoV-2 would become infected by the virus (the “probability of susceptibility”).

### Wildlife Rehabilitators Infection Risk Model

The WR infection risk model was calculated from two encounter types: bat handling, and workers in proximity to bats but not in a shared enclosed space. The expected number of infected bats arising from wildlife rehabilitation over the summer season is the sum of the expected number of bats infected through each of the two encounter types:

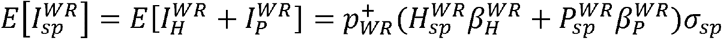

where

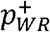 is the probability that someone conducting rehabilitation work is actively shedding SARS-CoV-2 virus on any given day of the 2021 active season;
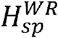 is the total number of bats handled during the 2021 active season;
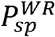 is the total number of bats exposed, not in an enclosed space or handled, by wildlife rehabilitators during the 2021 active season;
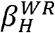 is the probability that a bat handled by a WR who was actively shedding virus would be exposed to the virus (an “exposure probability”) in the absence of any new restrictions, regulations, or protocols, taking into account the handling time typical of rehab activities;
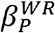 is the probability that a bat **not** in an enclosed space within a 6-foot proximity of (and not handled by) a WR who was actively shedding virus would be exposed to the virus (an “exposure probability”) in the absence of any new restrictions, regulations, or protocols; and
σ_*sp*_ is the species-specific probability that a bat exposed to a sufficient viral dose of SARS-CoV-2 would become infected by the virus (the “probability of susceptibility”).

### Wildlife Control Operators Infection Risk Model

The WC infection risk model is calculated from two encounter types: bat handling, and workers in proximity to bats but not in a shared enclosed space. The expected number of infected bats arising from wildlife control operations over the summer season is the sum of the expected number of bats infected through each of the two encounter types:

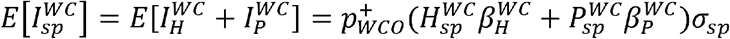

where

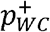 is the probability that someone conducting WC work is actively shedding SARS-CoV-2 virus on any given day of the 2021 active season;
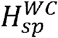 is the total number of bats handled during the 2021 active season;
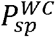 is the total number of bats exposed, but not handled, by WC during the 2021 active season;
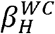 is the probability that a bat handled by a WC who was actively shedding virus would be exposed to the virus (an “exposure probability”) in the absence of any new restrictions, regulations, or protocols, taking into account the handling time typical of WC activities;
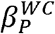 is the probability that a bat **not** in an enclosed space within a 6-foot proximity of (and not handled by) a WC who was actively shedding virus would be exposed to the virus (an “exposure probability”) in the absence of any new restrictions, regulations, or protocols; and
σ_*sp*_ is the species-specific probability that a bat exposed to a sufficient viral dose of SARS-CoV-2 would become infected by the virus (the “probability of susceptibility”).

### Probability that a crew member is positive and shedding virus

We calculated the probability that a crew member is positive and shedding virus as a function of the prevalence of COVID-19 in the surrounding community (*ψ*), and the sensitivity (*Sn*) and specificity (*Sp*) of COVID-19 testing. Sensitivity is the probability that an individual who has COVID-19 tests positive, whereas specificity is the probability that a healthy individual without COVID-19 tests negative. We selected a sensitivity value of 0.70, and specificity of 0.95 (Arevalo-Rodriguez et al. 2020; Watson et al. 2020); however, we recognize that these values vary according to the type of test administered. For our risk assessment, we are primarily interested in the probability that a crew member receives a negative test result but is truly infected with SARS-CoV-2. This probability can be calculated using Bayes’ Theorem as:

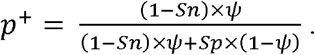

If a crew member does not take a test, the probability that a crew member is positive and shedding virus can be estimated by the local prevalence, *ψ*, or by some other method that accounts for the crew member’s risk behavior (e.g., https://www.microcovid.org/).

### Bat Encounter Types and Exposure Probabilities

To calculate the number of bats handled (*H*), encountered in an enclosed space (*E*), or in proximity to workers in an unenclosed space (*P*), we multiplied the total number of bats encountered in a typical season of work by the percentage of each bat encounter type (Table 1). We used the same encounter estimates reported in Runge et al. (2020), based on reporting data from Colorado Department of Wildlife, Connecticut Department of Energy and Environmental Protection, Kentucky Department of Fish and Wildlife Resources, New York State Department of Environmental Conservation, Oregon Department of Fish and Wildlife, Virginia Department of Game and Inland Fisheries, Wisconsin Department of Natural Resources, USDA Forest Service, National Park Service, U.S. Geological Survey, and the white-nose syndrome national surveillance program. For RSM activities, most bat interactions involve handling (45.8%), followed by activities in proximity to bats (42.7%), and sharing an enclosed space with bats (11.5%). For WR, all documented human bat interactions result from handling (100%). For WC, most human-bat interactions occur when a practitioner comes within 6-feet of a bat (77.1%) but does not handle the bat; the other 22.9% of interactions involve bat handling (Table 1).

**Table 1.**
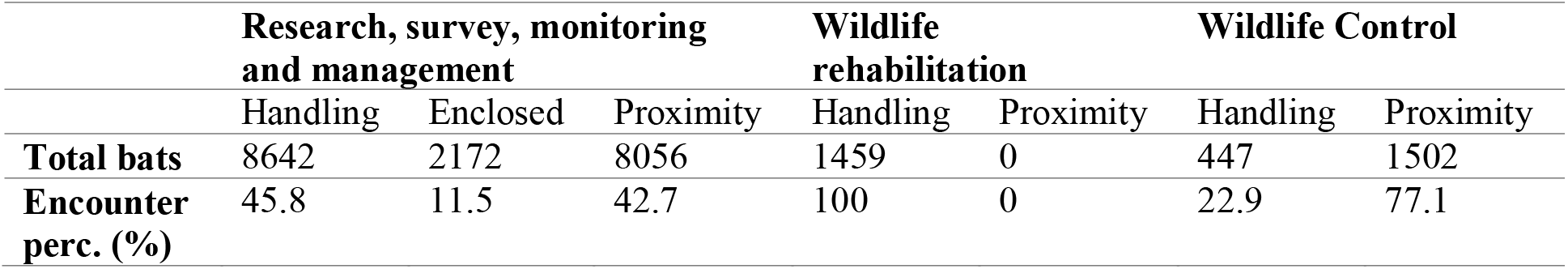
Fraction of bats encountered through each of three encounter types for each of three activities. The three encounter types are: handling; encounter within 6 feet in an enclosed space; and proximity within 6 feet in an unenclosed space. The data were gathered from the following agencies: Colorado Department of Wildlife, Connecticut Department of Energy and Environmental Protection, Kentucky Department of Fish and Wildlife Resources, New York State Department of Environmental Conservation, Oregon Department of Fish and Wildlife, Virginia Department of Game and Inland Fisheries, Wisconsin Department of Natural Resources, Forest Service, National Park Service, U.S. Geological Survey, and the White-Nose surveillance program (Runge et al. 2020).

|  | **Research, survey, monitoring**<br>**and management** |  |  | **Wildlife**<br>**rehabilitation** |  | **Wildlife Control** |  |
| --- | --- | --- | --- | --- | --- | --- | --- |
|  | Handling | Enclosed | Proximity | Handling | Proximity | Handling | Proximity |
| **Total bats** | 8642 | 2172 | 8056 | 1459 | 0 | 447 | 1502 |
| **Encounter**<br>**perc. (%)** | 45.8 | 11.5 | 42.7 | 100 | 0 | 22.9 | 77.1 |

<sup>1</sup>

For each human-bat activity and encounter type, Runge et al. (2020) used formal expert judgment protocols, notably the IDEA(“Investigate, Discuss, Estimate, Aggregate”) protocol (Hanea et al. 2017) and the four-point elicitation method (Speirs-Bridge et al. 2010), to estimate unique probabilities of exposure and an associated measure of uncertainty (i.e., β parameters in infection risk model equations). The four-point elicitation method provided a point estimate and a measure of within-expert uncertainty by eliciting each expert’s lowest, highest, and best estimates of model parameters as well as an estimate of confidence that their reported values included the true value. The expert panel included a total of 13 individuals with diverse professional experience and expert specializations in wildlife epidemiology, virology, bat physiology, and bat ecology (Runge et al. 2020). Two rounds of elicitation were held, and group meetings in between rounds were used to clarify questions and responses, with the aim of reducing sources of expert bias. To estimate an aggregate expert distribution from individual responses, the parameters that best fit probability distributions to the elicited quantiles from each expert independently were identified. Then parameters for an aggregate distribution were estimated by averaging the independent probability density functions (PDFs) across all experts and finding parameters for a fitted aggregate distribution that minimized the Kullback-Leibler distance (Kullback and Leibler 1951) between the average PDF and the fitted PDF.

For RSM activities, experts estimated a median of 49.7 bats (80% confidence interval: 15.3, 84.3) exposed out of 100 encountered during handling, a median of 19.4 bats (80% CI: 2.2.,72.4) exposed out of 100 encountered when in enclosed space within 6 feet of a SARS-CoV-2 positive scientist, and a median of 6.4 bats (80% CI: 0.6, 43.8) exposed out of 100 within 6 feet of a SARS-CoV-2 positive scientist in an unenclosed space. For WR activities, experts estimated a median of 70.4 bats (80% CI: 24.4, 94.6) exposed out of 100 during handling and a median of 24.3 bats (80% CI: 2.8, 78.4) exposed out of 100 when within 6 feet of a SARS-CoV-2 positive wildlife rehabilitator. For WC activities, experts estimated a median of 27.7 bats (80% CI: 3.7, 79.2) exposed out of 100 during handling and 9.6 bats (80% CI: 1.0, 53.9) exposed out of 100 when within 6 feet of a SARS-CoV-2 positive wildlife control operator.

In addition to COVID-19 testing for risk mitigation, agencies can issue guidelines for properly fitted PPE use during human-bat interactions. In Runge et al. (2020), the effectiveness of N95 respirators for mitigating the risk of SARS-CoV-2 exposure during RSM, WR, and WC activities was evaluated. Following that publication, additional information identified aerosolized virus as the primary pathway of human-to-human disease exposure (Meyerowitz et al. 2020); thus, we can evaluate the ability of other face coverings to reduce viral exposure of bats if we can assume that exposure probabilities from Runge et al. (2020) are reduced by reported filtration efficiencies of other PPE types. Common PPE types include: N95 respirators (percent filtration efficiency (FE) mean ± SD: 99.4 ± 0.2; 3M model 1870), surgical masks (FE: 89.5 ± 2.7), cloth masks (FE: 50.9 ± 16.8), and face shields (FE: 23 ±3.3) (Davies et al. 2013, Lindsley et al. 2014, Long et al. 2020).

### Probability of Susceptibility

We used probability of bat susceptibility (σ_*sp*_) estimates that were derived from expert judgment using the same structured protocols described above. In the Cook et al. (2021) application, the expert panel included a diverse group of 12 professionals, 4 of whom participated in the Runge et al. (2020) study. Similar to the Runge et al. (2020) study, two rounds of elicitation were held, and the probability of susceptibility for little brown bats was estimated using fitted aggregate group distributions based on the 12 expert responses.

The probability-of-susceptibility estimates, probability of infectious crew members, and effectiveness of PPE were then used to estimate the number of little brown bats that could be infected out of 1000 encountered during RSM, WC, and WR activities. For all comparisons, unless otherwise specified, we assumed that the local COVID-19 prevalence was 0.05. Each infection risk model was simulated 100,000 times to explore uncertainty in the parameters. All analyses were performed in Program R (R Core Team, 2018). All data used in these analyses were provided as electronic records and no vertebrate species were contacted or handled as a direct result of this study.

## RESULTS

### Probability of Susceptibility

Conditional on a sufficient dose of SARS-CoV-2 for individual bat infection, the expert panel from Runge et al. (2020) estimated that the median probability of susceptibility for little brown bat was 0.44 (80% PI:0.08 – 0.88). Following the accumulation of new information, a follow-up expert elicitation estimated that the median probability of susceptibility was 89% lower and had less uncertainty (Cook et al. 2021; Figure 2, median probability of susceptibility: 0.05; 80% PI:0.003 – 0.37). The updated estimate was informed, in part, by new information, including human and bat angiotensin-converting enzyme 2 (ACE2) receptor homology (an enzyme that is important to SARS-CoV-2 binding in the host) (Damas et al. 2020), and the availability of lab-based challenge studies (Hall et al. 2020).

**Figure 2.**
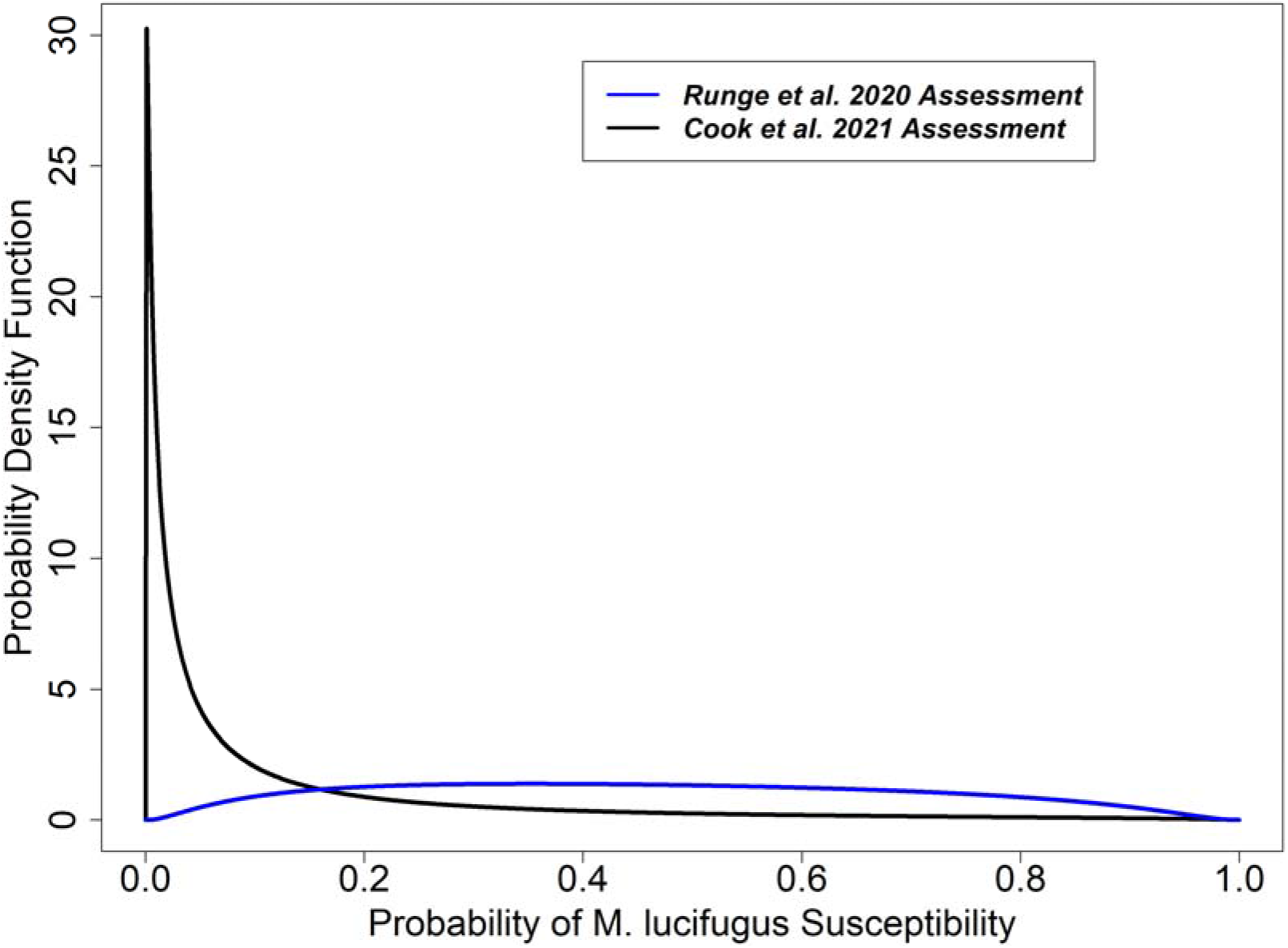
Comparison of probability of susceptibility estimates for little brown bat from Runge et al. (2020) (blue line) and from Cook et al. (2021) (black line).Experts estimated that the median probability of susceptibility was 89% lower based on updated knowledge gathered from bat challenge studies, ACE2 homology between humans and bats, and other sources (Cook et al. 2021).

### Baseline risk

We reanalyzed the infection risk models described in Runge et al. (2020) using updated estimates of the probability of susceptibility for little brown bats reported in Cook et al. (2021). We found an 87 – 88% decrease in the median number of bats estimated to be infected per 1000 encountered when compared against the earlier results. For RSM activities, the median number of bats infected per 1000 was estimated to be 6.96 in the Runge et al. (2020) assessment (Figure 3; 80% CI: 1.85 – 19.41). Using updated probability of susceptibility estimates, we found that the median number of bats estimated to be infected by SARS-CoV-2 was less than one individual per 1000, which is 88% lower than the initial estimate (Figure 3; median: 0.83, 80% CI: 0.07 – 7.82). For WR encounters, the median number of bats infected per 1000 was reduced from 13.03 (Figure 3; 80% CI: 3.54 – 36.14) to 1.56 – a similar 88% decrease in the median value (Figure 3; 80% CI: 0.12 – 14.71). For WC encounters, the median number of bats infected per 1000 was reduced from 3.72 (Figure 3; 80% CI: 0.84 – 14.43) to 0.47 – an 87% decrease in the median value (Figure 3; 80% CI: 0.03 – 4.79).

**Figure 3.**
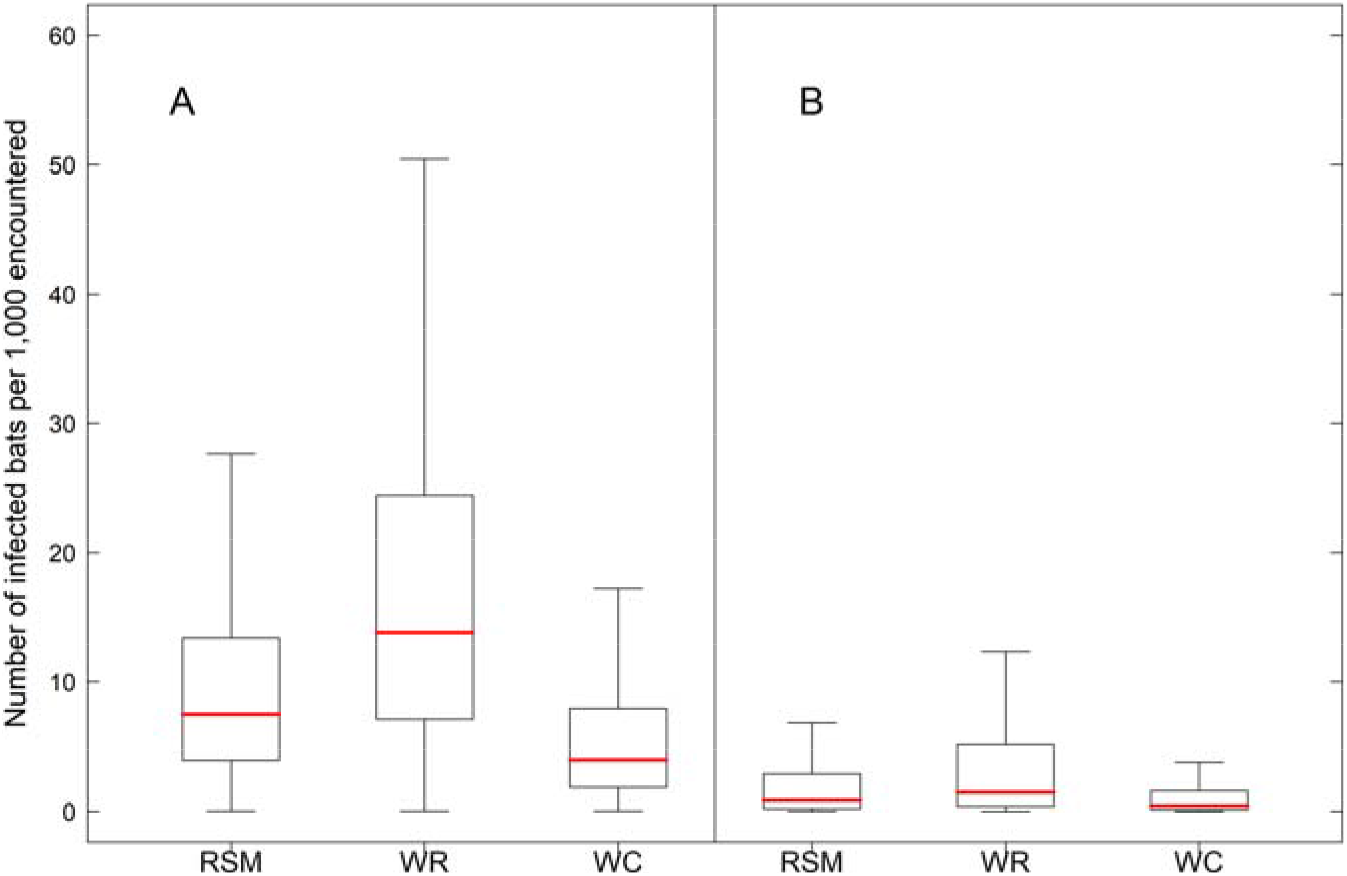
Number of bats per 1,000 exposed to and infected by SARS-CoV-2 by the three transmission pathways. RSM=research, survey, monitoring, and management activities; WR= wildlife rehabilitation; WC= wildlife control operations. Boxplot whiskers represent 99% prediction interval. For comparisons, we used the same assumed ratio of encounter modes (handling, enclosure, and proximity) and probability of worker shedding SARS-CoV-2 (median: 0.057; 80% interval: 0.022-0.112) from Runge et al. (2020). **(A)** Results reproduced based on expert-elicited data on probability of bat susceptibility from Runge et al. (2020) assessment. **(B)** Results based on parameter values from December 2020 assessment.

We also analyzed the baseline bat infection risk across three different levels of COVID-19 prevalence (Figure S.1). For RSM activities, the median number of bats infected per 1000 encountered fell from a median of 0.83 (80% CI: 0.07 – 7.82) when COVID-19 prevalence was 0.05, to a median of 0.15 (80% CI: 0.01 – 1.29) and 0.015 (80% CI: 0.001 – 0.129) when COVID-19 prevalence was 0.01 and 0.001, respectively. For WR activities, the median number of bats infected per 1000 fell from a median of 1.56 (80% CI: 0.12 – 14.71) when COVID-19 prevalence was 0.05, to a median of 0.27 (80% CI: 0.02 – 2.34) and 0.027 (80% CI: 0.002 – 0.23) when COVID-19 prevalence was 0.01 and 0.001, respectively. For WC activities, the median number of bats infected per 1000 fell from a median of 0.47 (80% CI: 0.03 – 4.79) when COVID-19 prevalence was 0.05, to a median of 0.08 (80% CI: 0.005 – 0.76) and 0.008 (80% CI: 0.0005 – 0.08) when COVID-19 prevalence was 0.01 and 0.001, respectively.

### Risk mitigation

We used the updated parameter estimates to evaluate the effectiveness of PPE and pre-survey COVID-19 testing of personnel for reducing baseline estimates of bat infections per 1000 encountered (Figures 4 and 5). We found that N95 respirators reduced the median estimates of infection by 95 – 96% for all three encounter types (i.e., RSM, WR, and WC work), when compared against median values with updated parameter estimates and without enhanced personal protective equipment (Figure 4A – C; overall reduction of 99% from Runge et al. 2020). For surgical masks, we found an 89% reduction in the median estimate of infection for all three encounter types (i.e., RSM, WR, and WC work; Figure 4A – C), when compared against median values with updated parameter estimates and without enhanced personal protective equipment. For cloth masks, we found a reduced median estimate of infection of 54 – 55% for all three encounter types (Figure 4A – C), when compared against median values with updated parameter estimates and without enhanced personal protective equipment. Finally, for face shields, we found a reduced median estimate of infection of 22 – 24% for all three encounter types (Figure 4A – C), when compared against median values with updated parameter estimates and without enhanced personal protective equipment.

**Figure 4.**
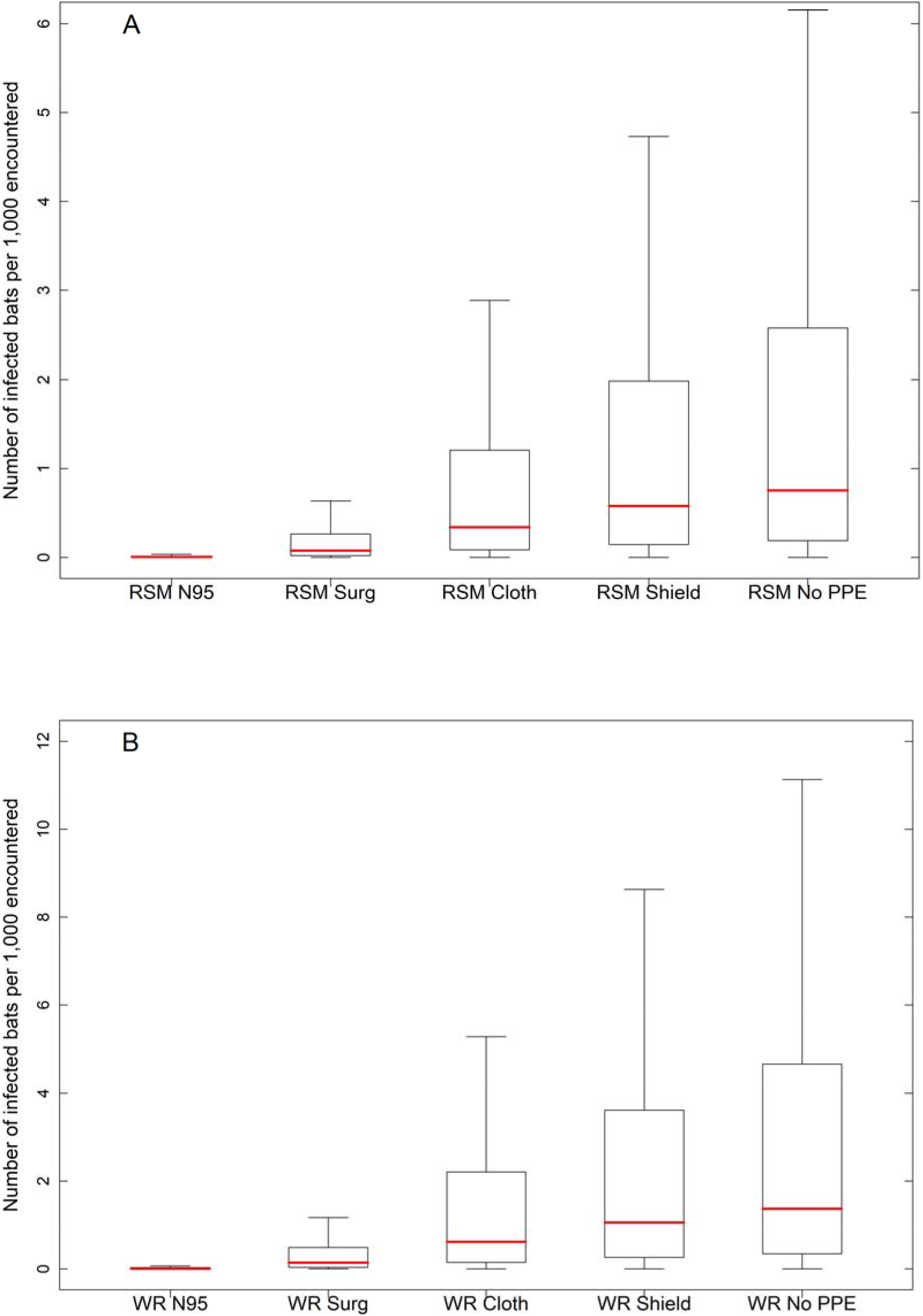
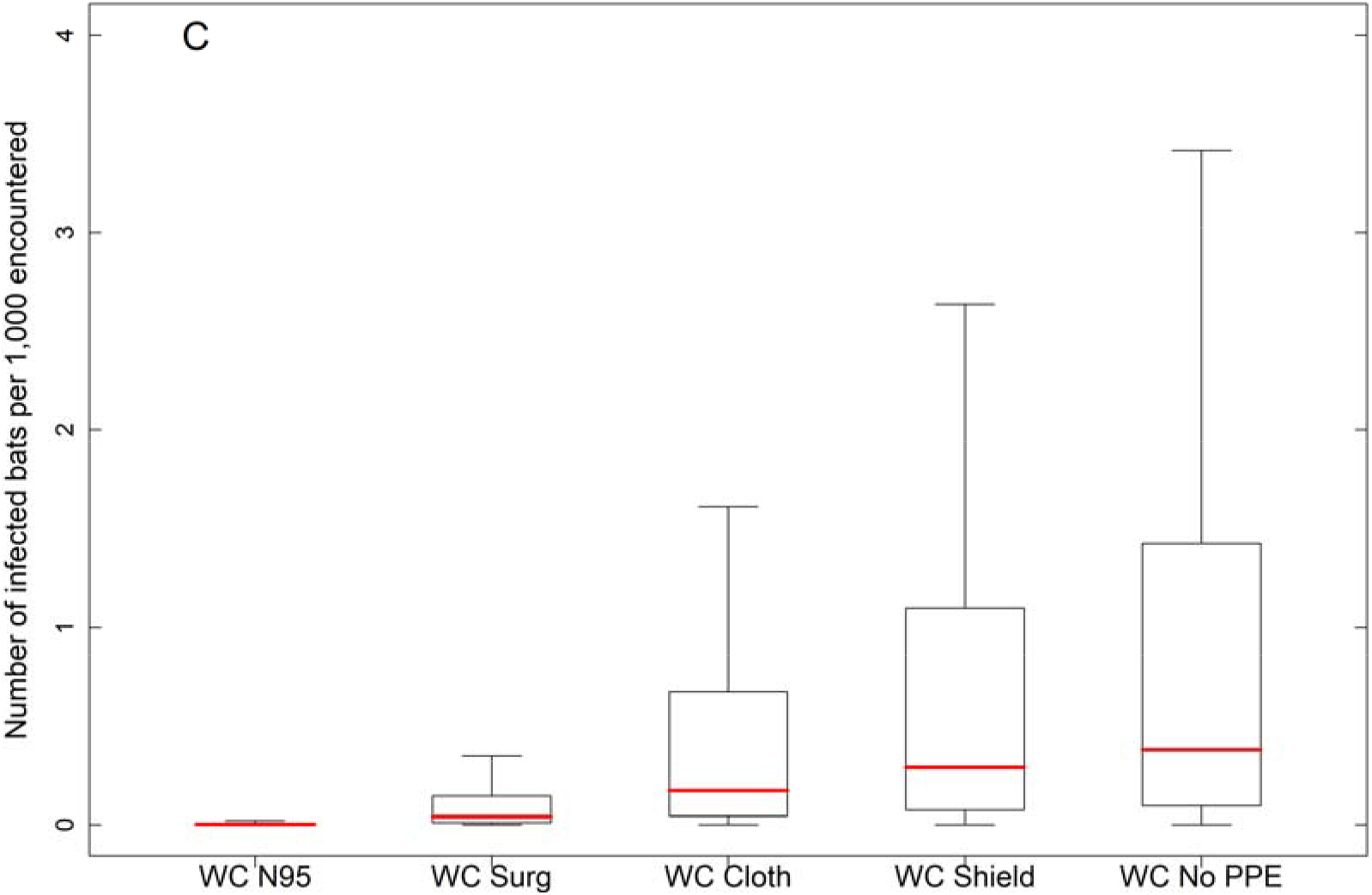
Number of bats per 1,000 exposed to and infected by SARS-CoV-2 by the three transmission pathways. RSM=research, survey, monitoring, and management activities; WR= wildlife rehabilitation; WC= wildlife control operations. Boxplot whiskers represent 99% prediction interval. We used the same assumed ratio of encounter modes (handling, enclosure, and proximity) from Runge et al. (2020). Results based on expert elicited data on probability of bat susceptibility from the Cook et al. (2021) assessment. **(A)** Effectiveness of PPE compared against baseline estimates for RSM activities. **(B)** Effectiveness of PPE compared against baseline estimates for WR activities. **(C)** Effectiveness of PPE compared against baseline estimates for WC activities.

**Figure 5.**
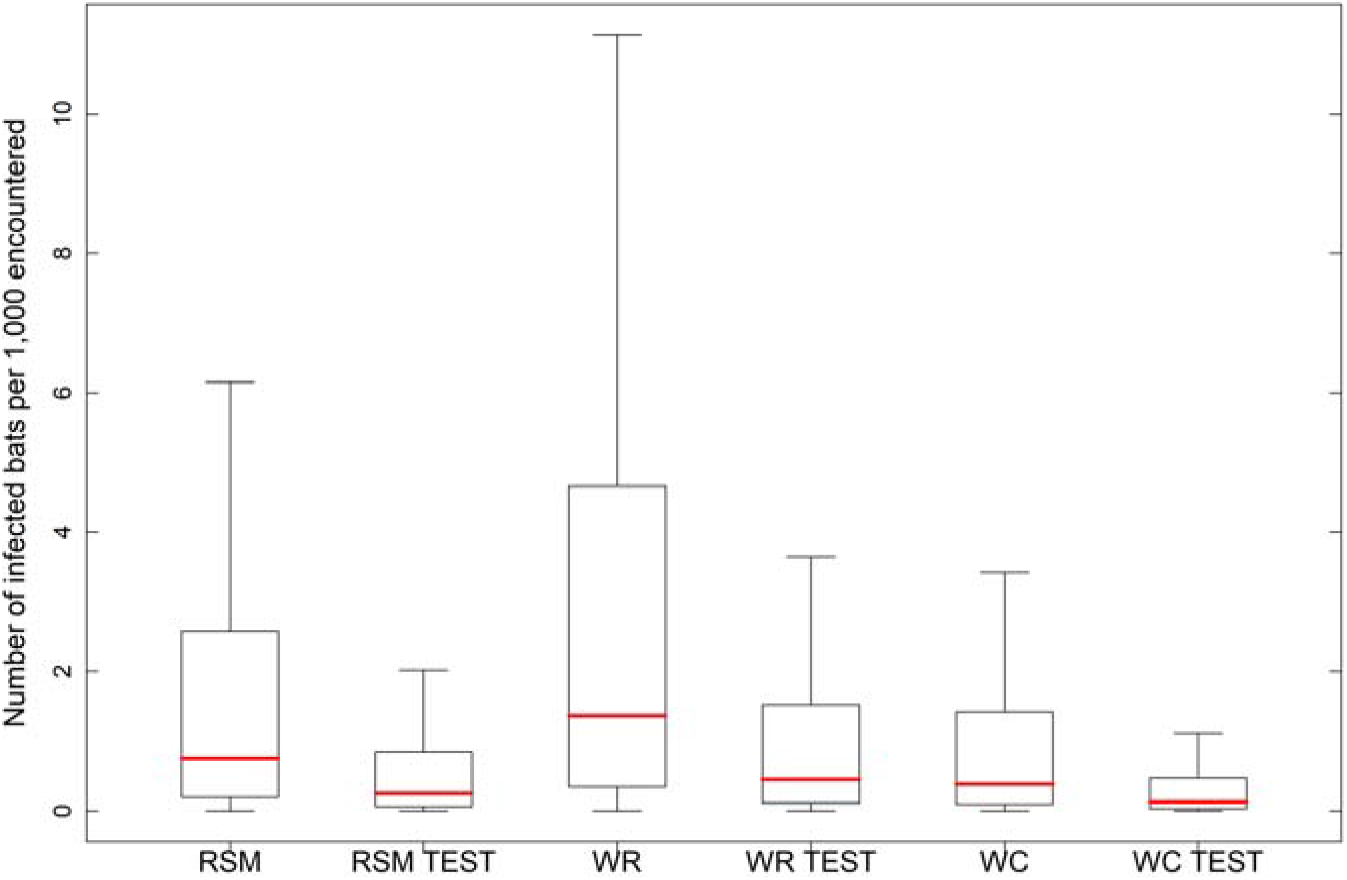
Number of bats per 1,000 exposed to and infected by SARS-CoV-2 by the three transmission pathways with and without pre-survey COVID-19 testing. RSM=research, survey, monitoring, and management activities; WR= wildlife rehabilitation; WC= wildlife control operations. Boxplot whiskers represent 99% prediction interval. We used the same assumed ratio of encounter modes (handling, enclosure, and proximity) from Runge et al. (2020). Results based on expert elicited data on probability of bat susceptibility from Cook et al. (2021) assessment.

For COVID-19 testing, we found that the median estimate of the number of SARS-CoV-2 infected bats decreased by 65 – 67% across all three encounter types, as a result of a negative test of field crew 3 days prior to bat handling (overall reduction of 88 – 89% from the Runge et al. 2020 assessment; Figure 5).

## DISCUSSION

The existing decision framework developed in Runge et al. (2020) allowed for a rapid re-evaluation of human-to-bat SARS-CoV-2 transmission risk during summer fieldwork based on new knowledge included in Cook et al. (2021), expert judgment, and other empirical studies. We found that new knowledge substantially reduced uncertainty, lowered risk estimates, and provided additional management alternatives that may be important to preventing SARS-CoV-2 infection in bats during RSM, WC, and WR activities. More broadly, we found that decision analysis coupled with expert judgment provided substantial benefits across all three studies (i.e., Runge et al. 2020, Cook et al. 2021, this study) and we expect that these benefits transcend SARS-CoV-2 to other wildlife disease systems. In particular, decision analysis helped to identify the fundamental management objectives, specify the possible alternatives, direct the development of quantitative infection risk models, and ultimately, create a risk assessment framework that remained useful over time and as our knowledge of the novel pathogen system improved. Formal expert judgment allowed us to estimate parameters with the available information in a timely manner, without having to initiate and wait for the results of new empirical studies.

We found that the median numbers of little brown bats potentially infected during summer RSM, WC, and WR activities were reduced substantially from those reported in the initial risk assessment by Runge et al. (2020), in part because an expanded range of management alternatives were available to further reduce these risks. By expanding the range of management alternatives for preventing transmission, we provide decision makers with additional options (and estimates of their effect on the number of infected bats) that may allow for some research and management activities to resume. For example, if the baseline risk of an activity exceeds an agency’s tolerance for risk, they may choose to require the use of facemasks or COVID-19 testing prior to human-bat encounters. However, COVID-19 testing may be difficult to implement in certain situations, especially for WC and WR activities that often arise spontaneously rather than from advanced planning. As an alternative, other forms of mitigation may be called for, and our assessment provides measures of risk across a range of potentially suitable mitigation measures. It is also important to note that risk tolerance may differ among agencies, and thus the response to the same risk may differ markedly in the decisions made (Sells et al. 2016).

Across the three risk assessments (Runge et al. 2020, Cook et al. 2021, this study), expert judgment was critical to our ability to rapidly estimate risks associated with SARS-CoV-2. At the time of Runge et al. (2020) and Cook et al. (2021), there were no data from empirical studies available to directly inform little brown bat SARS-CoV-2 susceptibility. Instead, structured protocols were implemented that derived unknown parameter estimates using leading experts in relevant fields of study. Expert judgment has gained credibility across a diversity of decision-making applications because it provides a viable alternative when empirical data are not yet available for generating unknown parameter estimates (Tyshenko et al. 2016; Bianchini et al. 2020), quantifying uncertainty (McBride et al. 2012; Conroy and Peterson 2013), and controlling for sources of bias (McBride et al. 2012). We found expert elicitation to be particularly powerful for our assessments because it allowed us to rapidly integrate the best available science and knowledge and provide guidance to managers dealing with uncertain but immediate risks to North American bats.

While decision framing, expert judgment, and the development of a quantitative infection risk model assisted in the production of decision-relevant science, the timely release of our results to support decision-making remained a challenge. Across the first two studies, the production of science happened over the course of several weeks, including several rounds of agency consultation and the development of quantitative infection risk models. For information sharing, we were able to provide the results to decision makers in a timely fashion through briefings after we had peer review and agency clearance, but before the results were published. The agencies, however, were also interested in timely publication, so they could cite the research when communicating their decisions to the public. In total, the documentation, external peer-review, and publication process added an additional 5.5 weeks for Runge et al. (2020) and 11.5 weeks for Cook et al. (2021). While these timelines are typical and generally necessary for rigorous peer review, shorter timelines for studies evaluating emergent wildlife disease risks could be helpful because the decisions they are intended to inform may be necessary before completion of a standard peer-review and publication process. While it is not our intention to criticize any journal, reviewers, or peer-review process, we recognize that the production of decision-relevant science using decision analysis, quantitative modeling, and undergoing a full peer-review process may benefit from shorter timelines to provide information needed for urgent agency decisions.

There are likely many options to improve the timely delivery of science to support urgent wildlife disease management decisions moving forward, and we provide a few suggestions that may be useful. First, it may be useful for journals to consider creating alternate production tracks that can expedite the review and publication process and provide timely results at the speed of agency decisions. Alternative options for distribution, such as preprint servers (like bioRxiv and medRxiv) have already become critical avenues for timely release of information during the COVID-19 pandemic; however, these avenues do not address the critical role that peer-review plays in the production of reliable science. Second, for agencies that frequently make urgent decisions and that currently rely only on published results to communicate scientific support for those decisions to the public, it may be beneficial to consider using external science review boards that can provide objective evaluations of unpublished findings for formal consideration in time-sensitive decision-making. Lastly, dedicated risk assessment teams that produce rapid qualitative assessments of wildlife disease risks within hours or days of an identified novel hazard would be helpful. While other, more qualitative, assessments may be based on preliminary results and limited knowledge that is subject to considerable change, they can be effective as a bridge to more rigorous assessments that include agency consultation, quantitative modeling, and the evaluation of management alternatives. Nevertheless, we hope that our risk assessments may serve as a model to assess threats that SARS-CoV-2 continues to present to wildlife, and that a larger discussion be stimulated to identify the best approaches to deliver decision-relevant science for emerging wildlife diseases on timescales that matter.

Moving forward, it will be possible to update future SARS-CoV-2 risk assessments as knowledge of the critical uncertainties improves. Currently there are several variants of the SARS-CoV-2 pathogen circulating in the human population that have the potential to alter the transmissibility of the virus for humans and may affect the susceptibility of wildlife species to the virus. There are widespread vaccination efforts occurring that will also reduce the localized risk that workers are infectious at the time of bat encounters. At the local level, vaccinations are likely to decrease the prevalence of COVID-19, ultimately reducing SARS-CoV-2 transmission risk to bats (Figure S.1). At the individual level, COVID-19 vaccines are effective at preventing human infection (median: 66.3% [95% confidence interval (CI): 59.9%–71.8%] effective for preventing symptomatic, lab confirmed cases 14 days post-immunization for Johnson & Johnson vaccine [Oliver et al. 2021], and median: 90% [95% CI: 68%–97%] effective for preventing all cases regardless of symptoms 14 days post-immunization for Pfizer mRNA vaccine [Thompson et al. 2021]); however, there remain unknowns surrounding the period of immunity and its efficacy against newly emerging viral strains. More information should be available in coming months on the duration of protection from vaccine and the potential for individuals with “breakthrough” infections to shed virus. As our knowledge continues to improve surrounding SARS-CoV-2 and the risks it presents to bats, these factors, as well as other relevant information, could be included in future assessments to ensure that agencies have the best available information for making decisions.

## ACKNOWLEDGMENTS

The manuscript was improved with comments from reviewers. Any use of trade, product, or firm names is for descriptive purposes only and does not imply endorsement by the U.S. Government.

## Supplementary Figure

**Figure S.1.**
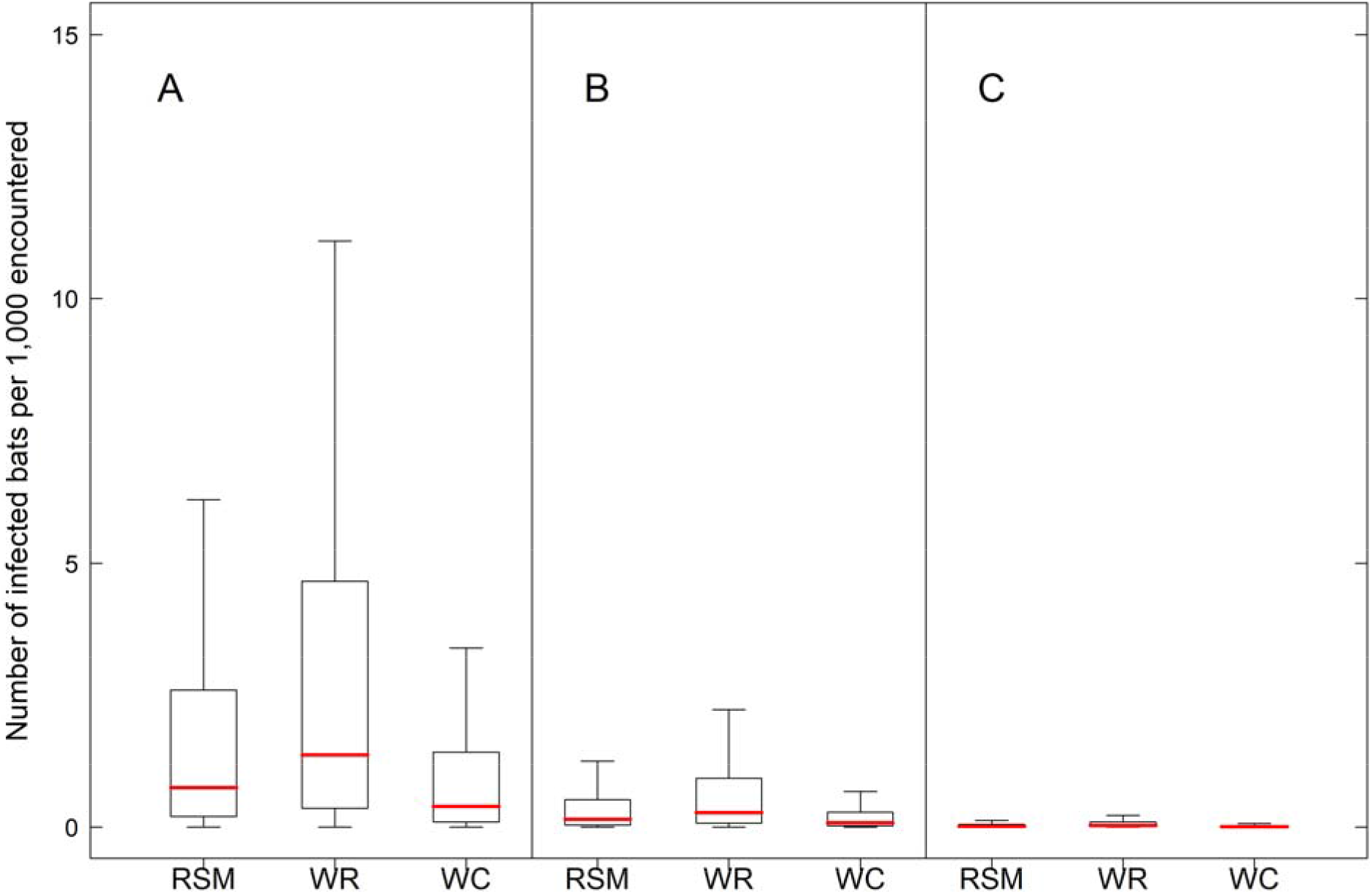
Comparison of the number of bats per 1,000 exposed to and infected by SARS-CoV-2 by the three transmission pathways and across 3 community prevalence levels: 0.05, 0.01, and 0.001. RSM, research, survey, monitoring, and management activities; WR, wildlife rehabilitation; WC, wildlife control operations. Boxplot whiskers represent 99% prediction intervals. We used the same assumed ratio of encounter modes (handling, enclosure, and proximity) from Runge et al. (2020). Results based on expert elicited data on probability of bat susceptibility from Cook et al. (2021) assessment. **(A)** Number of bats infected at county-level COVID-19 prevalence of 0.05. **(B)** Number of bats infected at county-level COVID-19 prevalence of 0.01. **(C)** Number of bats infected at county-level COVID-19 prevalence of 0.001.

